# Pyrethroid resistance intensity in *Anopheles gambiae* s.l. mosquito populations from Rwanda

**DOI:** 10.1101/2025.05.21.655281

**Authors:** Dunia Munyakanage, Lambert Nzungize, Torbert Yvan Migambi, Pascal Mosengo, Gerald Habimana, Aimable Mbituyumuremyi, Alphonse Mutabazi, Emmanuel Hakizimana

## Abstract

Evidence based vector control interventions depends on understanding the distribution and the evolution of insecticide resistance in malaria vectors. This study aims to assess the status of insecticide resistances in *Anopheles gambiae* s.l., to pyrethroid insecticides such as permethrin, deltamethrin and alphacypermethrin from 18 study sites located in 15 districts across Rwanda, representing the key strata of malaria transmission in country. The larvae of anopheles were collected from October 2022 to April 2023 using dipping method, and reared to adult stage. The *An. gambiae* were exposed to alphacypermethrin 0.05% (Acyp_1X), permethrin 0.75% (Perm_1X), and deltamethrin 0.05% (Delth_1X) then tested using WHO bioassay standard protocol for insecticide resistance monitoring. As results, we found an extremely high resistance to permethrin in Gashora (Bugesera district), Mubuga (Karongi district), Rwaza (Musanze district), Kirarambogo (Gisagara district) with mortality rate of 42%, 48%, 69% and 70% repsectively. Subsquently, no recovery of susceptibility was observed in permethrin after pre-exposure to piperonyl butoxide (PBO) and observed mortality was 70% in Kirarambogo, while in Gashora, Mubuga and Rwanza it was 80% at each site. In city of Kigali, the mortality with alphacypermethrin was 94% and 93% with synergist PBO. Notably, in Kicukiro district we found that metabolic resistance mechanism was not driving the resistance mechanisms. These findings provide important understandings of pyrethroid resistance status in Rwanda and offering valuable insights for distribution of insecticide resistance intensity across country. This highlights the need for further studies to assess the spread of insecticide resistance and molecular driven resistance mechanism to address the issue in malaria vector control in Rwanda.

## Introduction

In Rwanda two core vector control interventions have been implemented such as Indoor Residual Spraying (IRS) and Long-Lasting Insecticidal Nets (LLINs). Indoor Residual Spraying (IRS) was initially implemented in Kigali City’s three districts in 2007, then expanded to include rural districts with a significant malaria burden by 2011. The most commonly used class of insecticides for controlling malaria vectors, known as pyrethroids is becoming less effective against mosquito malaria vector species [1]. Currently, various classes of insecticides use in public health programs such as pyrethroids, organophosphates, carbamates, organochlorines, neonicotinoids, butenolides and butenolides etc. All are recommended by the World Health Organization Pesticide Evaluation Scheme (WHOPES) [2]. The pyrethroid class of insecticide is the mainstay of malaria vector control efforts and different malaria vector species have different mechanisms of insecticide resistance [3]. Implementing effective resistance management techniques throughout Rwanda requires an understanding of malaria interventions strategies used in Rwanda [4], emergence and distribution of insecticide resistance in major malaria vectors, for instance *An. gambiae*. The complex of sibling species known as Anopheles gambiae (sensu lato) (s.l.) includes An. gambiae (s.s.) and An. arabiensis, which are the two primary malaria vectors in sub-Saharan Africa (SSA) and in Rwanda. *An. gambiae* is among the dominant anopheles species responsible for the transmission of malaria parasites in Africa [5]. A primary cause of pyrethroid resistance is the decreased sensitivity of the voltage-gated sodium channel (Vgsc), which is the target site. Knockdown resistance (kdr) refers to the group of mutations in the Vgsc gene that result in amino acid substitutions and confer pyrethroid resistance [6]. Currently, they are limited published information about the extent of pyrhtroid resistance status and its geographical distribution in Rwanda. Consequently, it remains patchy whether the malaria vector control strategy could efficiently control malaria vector species like *An. gambiae* population across Rwanda. Furthermore, the increased of insecticide resistance identified during nationawide survey in Rwanda between 2011 and 2013 [7], if it stills prevailing throughout country, it remains unknown whether resistance is driven by metabolic resistance mechanisms or not. To fill these knowledge gaps, in this study, we extensively determined the distribution of resistance status of *An. gambiae* population from Rwanda and assessed the potential restoration of susceptibility through synergists assay using PBO to several insecticides used in Rwanda such as permethrin (Type I), deltametrhin (Type II) and alphacypermthrin (Type II).

## 2. Material and methods

### 2.1 Mosquito collection

The larvae of anopheles mosquitoes were collected across country over the four provinces (Fig. 1) of Rwanda in two districts of Western province (Karongi district and Nyamasheke district), six districts in Eastern province (Bugesera district, Nyagatare district, Rwamagana district, Kirehe district, Kayonza district, and Ngoma district), four districts of Northern province (Burera district, Rulindo district, Musanze district, and Gicumbi district). Furthermore, we collected in two districts of Southern province (Ruhango district and Nyamagabe district) as well as one district in Kigali city (Kicukiro district). The mosquito larvae were collected from October 2022 to April 2023. We collected larvae using dipping method (350 ml) and kept in a tray bucket under a cool condition prior to transport to the insectary at Rwanda Biomedical Center where they were reared up to adult stage. Based on morphological identification of adult mosquitoes (Coetzee 2020) used for resistance tests, we identified *An. gambiae* s.l complex and An. funestus group.

**Figure 1.**
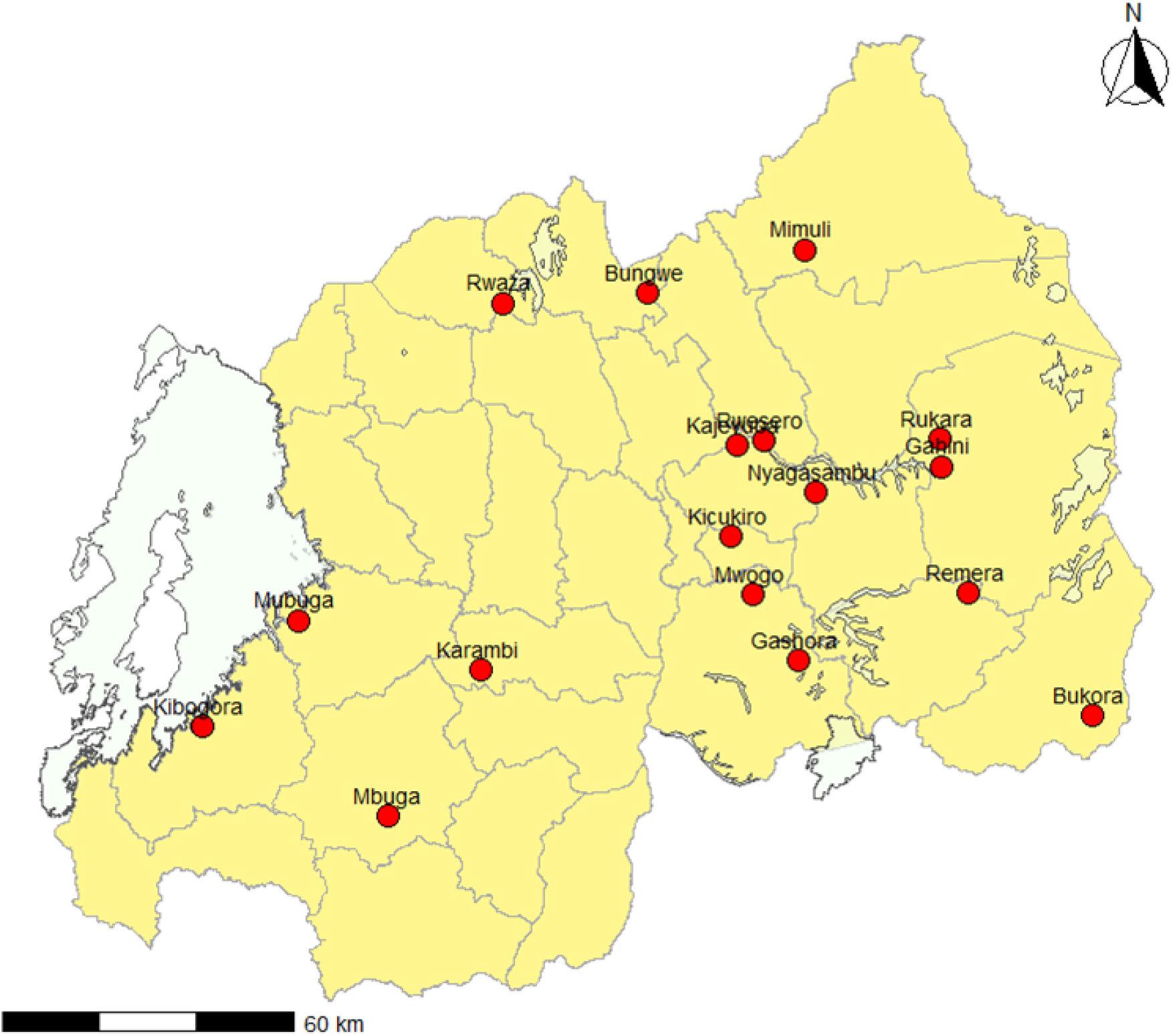
Collection sites of *An. gambiae* across the country

### 2.1 Insecticide susceptibility test

The susceptibiliy tests were performed on adult female mosquitoes using the WHO insecticide susceptibility tube-test procedures using standard diagnostic doses recommended for each insecticide [8]. For the bioassays to determine susceptibility, we used insecticide-impregnated filter papers such as permethrin (0.75%), deltamethrin (0.05%) and alphacypermethrin (0.05%). For the control, we used papers treated with silicone oil. The total of 150 female mosquitoes (2-4 day-old) were tested with four replicates each hold around 20– 25 mosquitoes per tube and two tubes without insecticide serving as negative control. The knockdown effect of insecticide was recorded after 15 min, 30 min and 60 min of exposure (Table 1-3). Therefore, after 60 min of exposure, all female mosquitoes were transferred to holding tubes, provided with 10% sugar water and placed under conditions of 20-26 °C and 80% RH. The mortality rates were recorded 24 hours post-exposure. After the bioassays, all mosquitoes were preserved individually in 1.5-ml Eppendorf tubes for further molecular analyses.

**Table 1.**
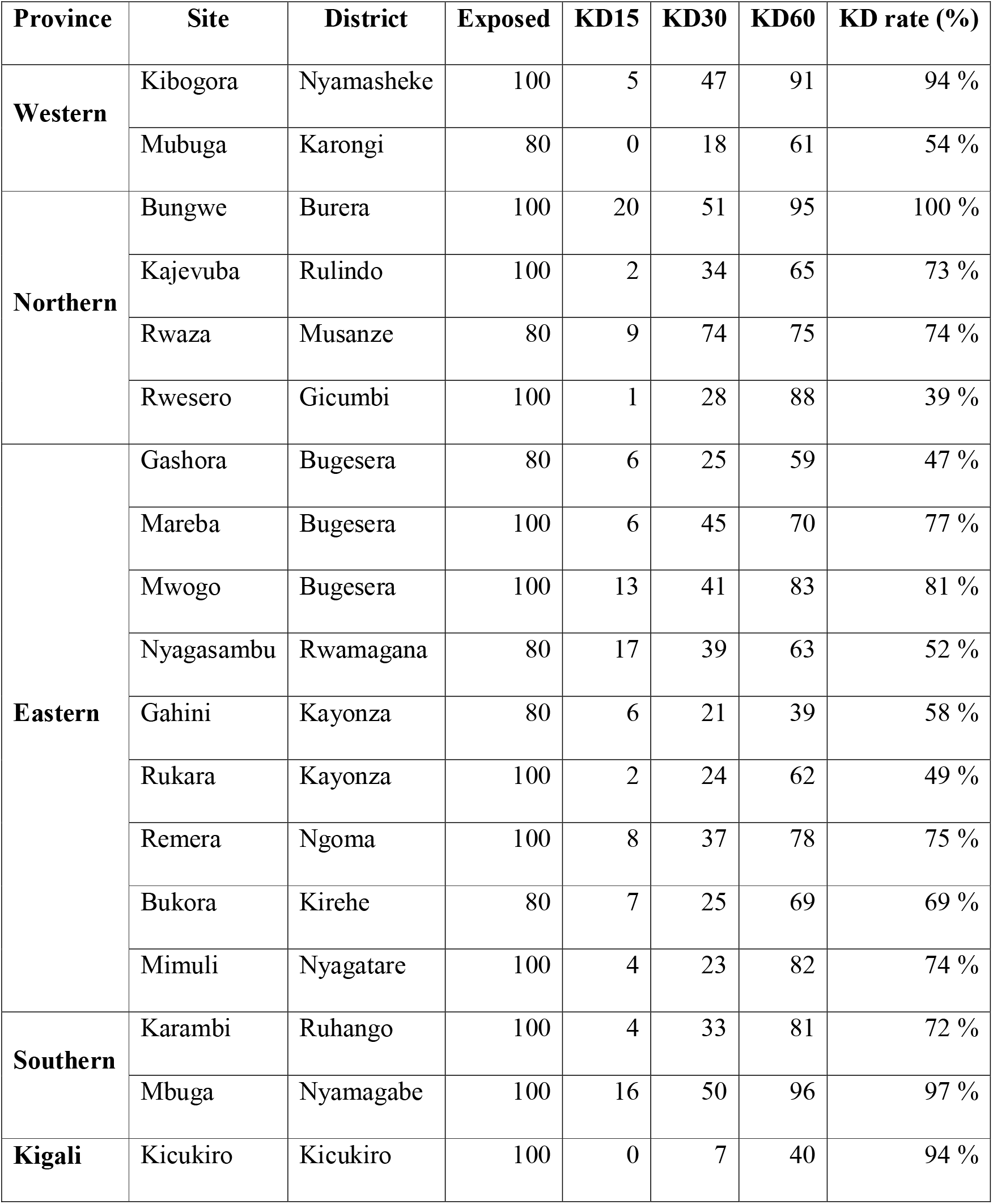
Knockdown effect of alphacypermethrin.

| Province | Site | District | Exposed | KD15 | KD30 | KD60 | KD rate (%) |
| --- | --- | --- | --- | --- | --- | --- | --- |
| **Western** | Kibogora | Nyamasheke | 100 | 5 | 47 | 91 | 94 % |
|  | Mubuga | Karongi | 80 | 0 | 18 | 61 | 54 % |
| **Northern** | Bungwe | Burera | 100 | 20 | 51 | 95 | 100 % |
|  | Kajevuba | Rulindo | 100 | 2 | 34 | 65 | 73 % |
|  | Rwaza | Musanze | 80 | 9 | 74 | 75 | 74 % |
|  | Rwesero | Gicumbi | 100 | 1 | 28 | 88 | 39 % |
| **Eastern** | Gashora | Bugesera | 80 | 6 | 25 | 59 | 47 % |
|  | Mareba | Bugesera | 100 | 6 | 45 | 70 | 77 % |
|  | Mwogo | Bugesera | 100 | 13 | 41 | 83 | 81 % |
|  | Nyagasambu | Rwamagana | 80 | 17 | 39 | 63 | 52 % |
|  | Gahini | Kayonza | 80 | 6 | 21 | 39 | 58 % |
|  | Rukara | Kayonza | 100 | 2 | 24 | 62 | 49 % |
|  | Remera | Ngoma | 100 | 8 | 37 | 78 | 75 % |
|  | Bukora | Kirehe | 80 | 7 | 25 | 69 | 69 % |
|  | Mimuli | Nyagatare | 100 | 4 | 23 | 82 | 74 % |
| **Southern** | Karambi | Ruhango | 100 | 4 | 33 | 81 | 72 % |
|  | Mbuga | Nyamagabe | 100 | 16 | 50 | 96 | 97 % |
| **Kigali** | Kicukiro | Kicukiro | 100 | 0 | 7 | 40 | 94 % |

### 2.2 Piperonyl butoxide (PBO) synergist assays

The adult mosquitoes were pre-exposed for one hour to piperonyl butoxide (4% PBO), then exposed to either alphacypermethrine, permethrin, or deltamethrin for another one hour. The mortality rate was assessed after 24 hours and compared with results obtained without pre-exposure to PBO to determine any possible involvement of cytochrome P450-mediated resistance.

#### Statistical analysis

Te mortality rates of the exposed mosquitoes and silicon oil as control samples were calculated per each insecticide used after bioassay. The mortality rate was determined using Abbott’s formula [9]. The status of mosquito resistance was interpreted based on the WHO guidelines as follows mortality rate < 90% its confrmed resistance, the mortality rate between 90-97% is possible resistance, and mortality rate ≥ 98% is susceptibility [8]. The data manipulation, statistical analyses and customize figures were performed using R packages such as ggpubr, dplyr, forcats and ggplot2. Results were represented based on standard errors (±SE) and p-value calculated based on Chi-square tests.

## 3. Results

### 3.1 Susceptibility profile to pyrethroids in Western province

We tested the insectice sucspecbility such as Permethrin, Deltamethrine and Alcypermthrin in two sites Kibogora located in Nyamasheke district and Mubuga located in Karongi district. We found that insecticide resistance was high in Nyamasheke district than karongi district where The mortality rate of *An. gambiae* to permethrin in Nyamasheke distric and Karongi district was 89% and 48% respectively. The mortality rate of *An. gambiae* to deltamethrin in Nyamasheke distric and Karongi district was 98% and 73% respectively. The mortality rate of *An. gambiae* exposed to alphacypermethrin in Nyamasheke distric and Karongi district was 94% and 54% respectively. We observed a partial recovery of susceptibility when mosquitoes were pre-exposed to piperonyl butoxide (Fig. 2)

**Figure 2.**
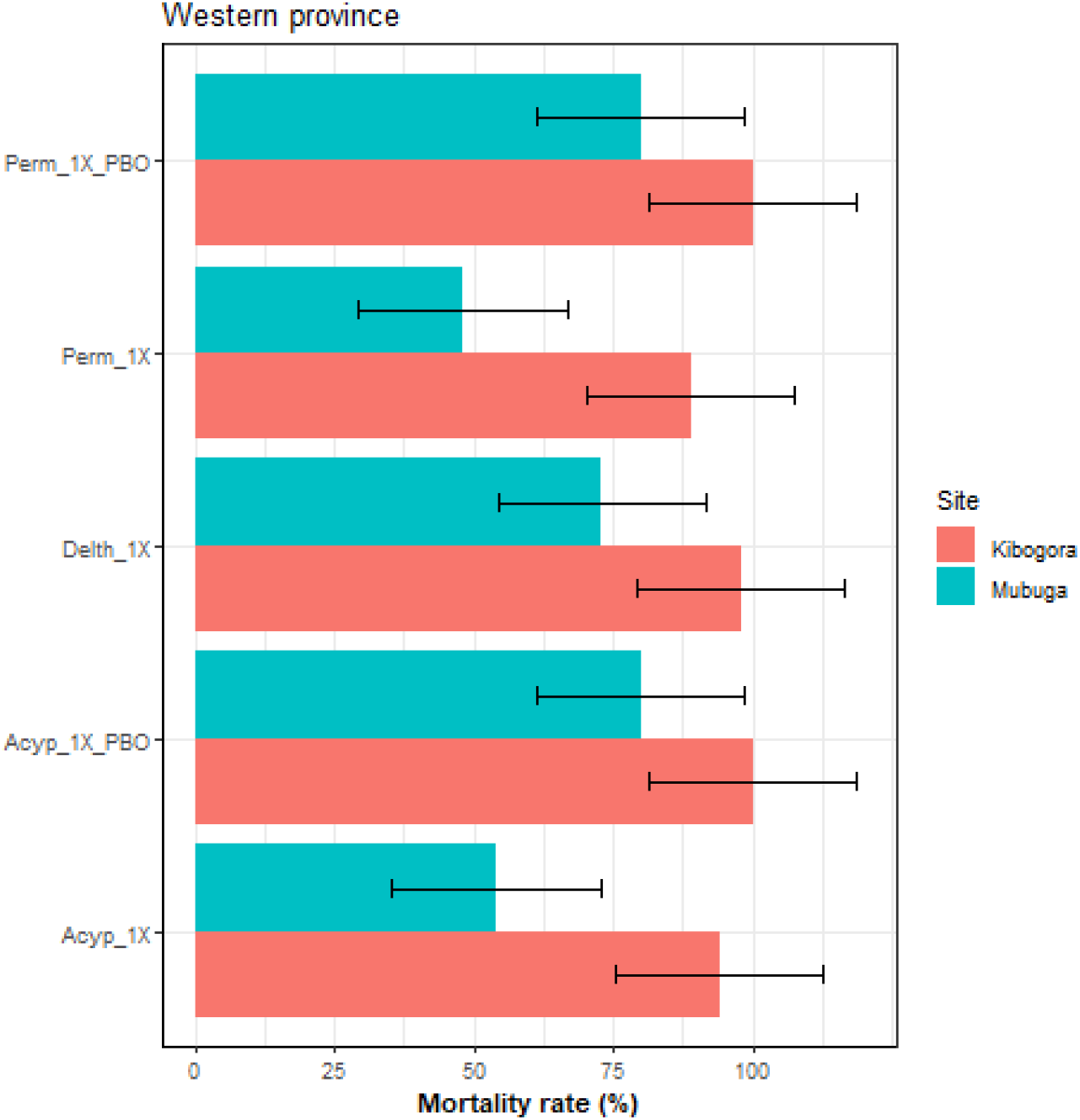
Susceptibility profile of *An. gambiae* population from Western province. Permethrin (Perm_1X), deltamethrin (Delth_1X) and alphacypermethrin (Acyp_1X). The effect of pre-exposure to synergist PBO against pyrethroids also represented (Perm_1X_PBO and Acyp_1X_PBO). The control had less than 5%. Results are the average of percentage mortalities and error bars represent ±SE of mean.

### 3.2 Susceptibility profile to pyrethroids in Eastern province

The bioassays using permethrin, alphacypermethrin and deltamethrin revealed a high resistance. The resistance against alphacypermethrin revealed that in Bugesera District the mortality rate was ranging from 47% to 81%, in Nyagatare district was between 58% and 74%. In other study sites from five districts such as Gatsibo, Ngoma, Kirehe, Rwamagana and Kayonza, the mortality recorded was 78%, 75%, 69%, 52%, 49% respectively. For permethrin resistance, the mortality observed in Bugesera district was ranging from 42% to 89%, in Kayonza district was 46% to 60%. Though in other districts such as Ngoma, Kirehe, Gatsibo, Rwamagana the mortality recorded was 79%, 68%, 57%, and 46% respectively. For deltamethrin profile, in Bugesera district mortality observed was ranging from 57% to 80%, in Kayonza district mortality was 55% to 78%. In other districts such as Gatsibo, Rwamagana, Kirehe, and Ngoma, the mortality rate recorded was 80%, 79%, 79%, and 55% respectively. For the mosquitoes pre-exposed to piperonyl butoxide (PBO), we observed a partial recovery of susceptibility (Fig. 3).

**Figure 3.**
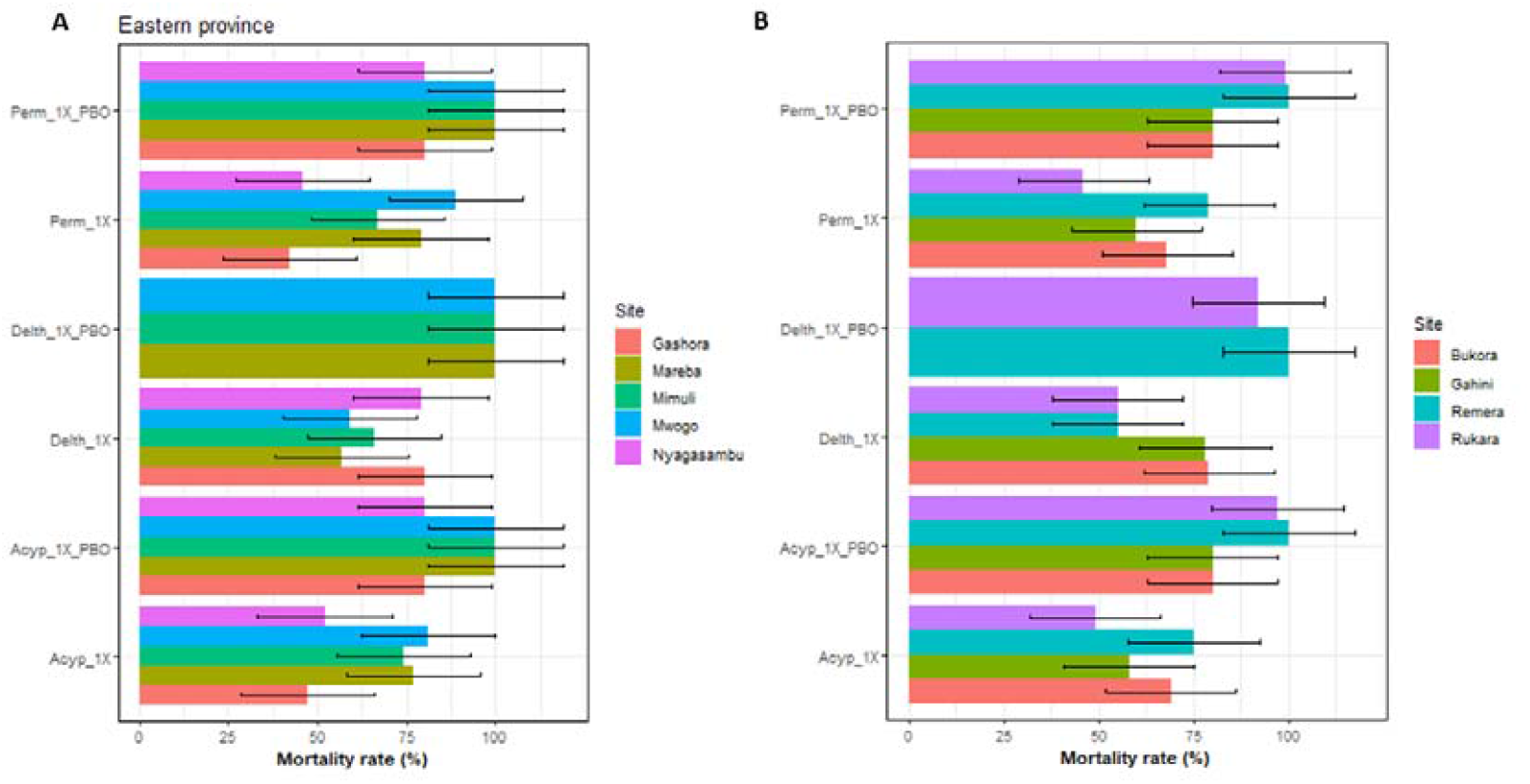
Susceptibility profile of *An. gambiae* population from Eastern province. Adult mosquitoes were exposed to permethrin (Perm_1X), deltamethrin (Delth_1X), and alphacypermethrin (Acyp_1X), both with and without pre-exposure to the synergist PBO(Perm_1X_PBO, Delth_1X_PBO, and Acyp_1X_PBO). Silicon oil was used as a control (≤ 5% mortality). Mortality was expressed as mean percentage ±SE.

### 3.3 Susceptibility profile to pyrethroids in Nothern province

We tested the level of resistance against pyrethroids such permethrin, deltamethrin and alphacypermethrin to mosquitoes population collected in Nothern province. For permethrin resistance, the mortality rate recorded in Burera district, Rulindo district, Musanze distric, and Gicumbi district was 100%, 71%, 69% and 38% respectively. We use bioassays to test alphacypermethrin gainst *An. gambiae* population collected in Burera district revealed a mortality rate of 100%, in Musanze district was 74%, Rulindo district was 73%, and Gicumbi distric was 39%. For deltamethrin against mosquitoes, the mortalite recorded in Burera district was 100%, in Rulindo district was 94%, in Musanze district was 78% and in Gicumbi district was 55% (Fig 4). A partial recovery of susceptibility was observed with the pre-exposed to piperonyl butoxide (PBO).

**Figure 4.**
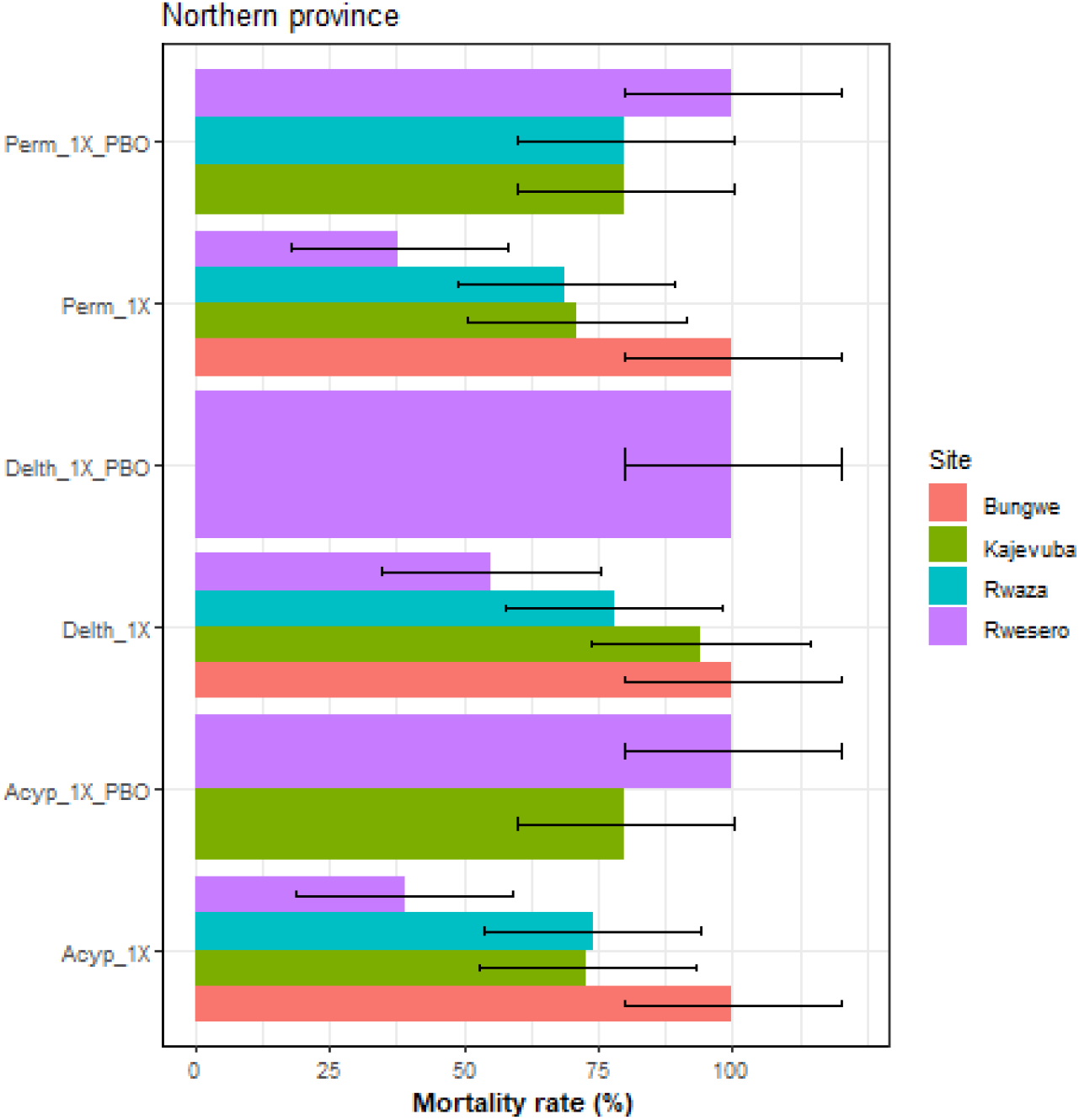
Susceptibility profile of *An. gambiae* population from Northern province. *An. gambiae* population were exposure to the three insecticides such as permethrin (Perm_1X), deltamethrin (Delth_1X) and alphacypermethrin (Acyp_1X). The effect of pre-exposure to synergist PBO against pyrethroids are also represented as (Perm_1X_PBO, Delth_1X_PBO and Acyp_1X_PBO). Silicon oil was used as a control (mortality ≤5%). Results are average of percentage mortalities and error bars represent ±SE of mean.

### 3.4 Susceptibility profile to pyrethroids in Southern province

The mortality observed for mosquitoe population exposed to permethrin from both districts susch as Nyamagabe was 97% and Ruhango was 52%. The mortality on exposure to deltamethrin in Nyamagabe district was 98% and in Ruhango district was 97%. For alphacypermethrin the mortality recorded in Nyamagabe and Ruhango districts was 97%, and 72% respectively. The pre-exposure of *An. gambiae* to PBO restored the susceptibility of mosquitoes population (Fig 5).

**Figure 5.**
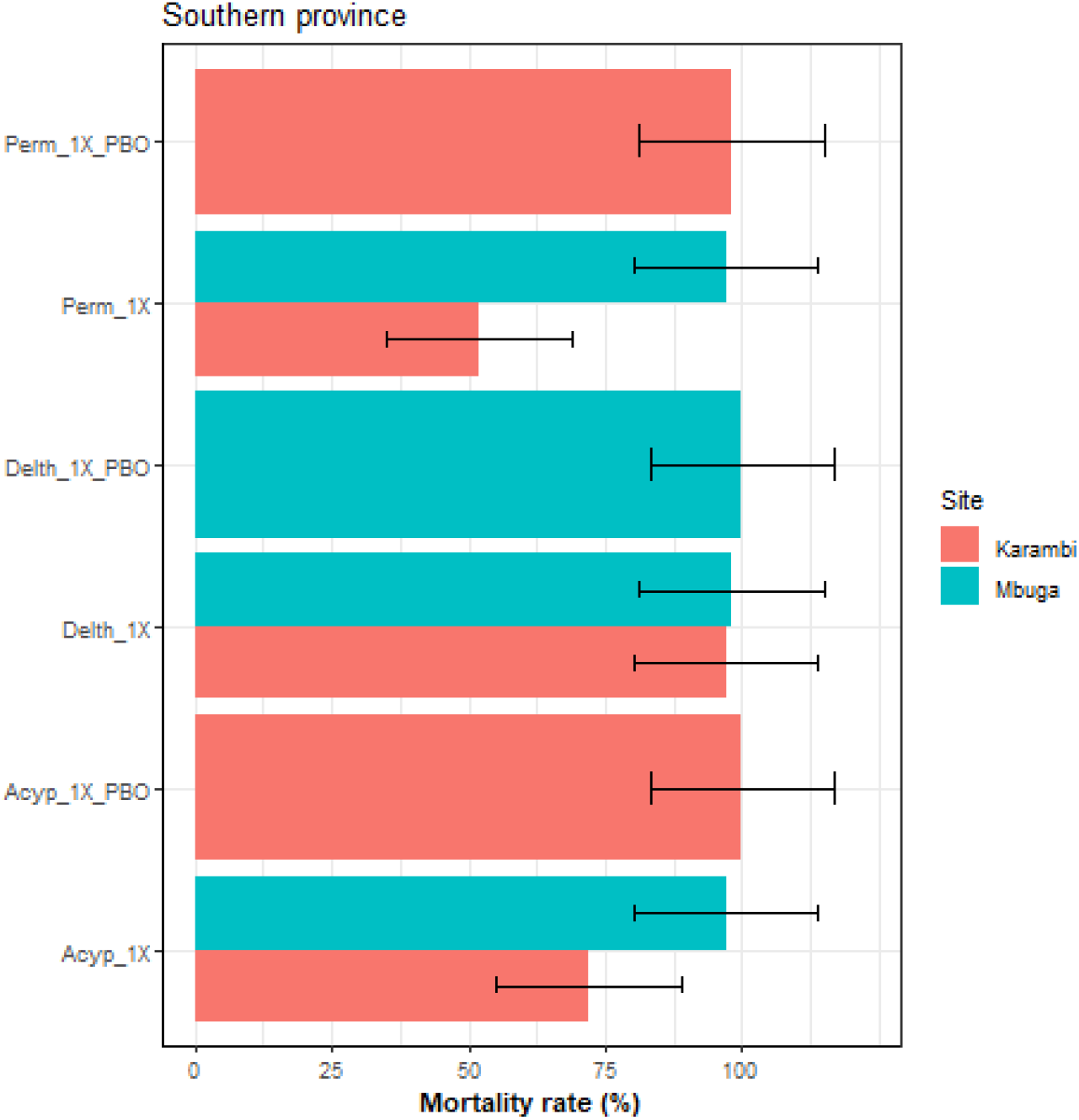
Susceptibility profile of *An. gambiae* population from Southern province. Three insecticides, such as deltamethrin (Delth_1X), alphacypermethrin (Acyp_1X), and permethrin (Perm_1X), were applied to the An. gambiae population. The effect of pre-exposure to synergist PBO against pyrethroids also represented by (Perm_1X_PBO, Delth_1X_PBO, and Acyp_1X_PBO). The control (silicon oil), had a mortality rate of less than 5%. Findings are the mean of the percentage mortality, and error bars represent ±SE of the mean.

### 3.5 Susceptibility profile to pyrethroids in the city of Kigali

The exposure for one hour of mosquitoes population collected from city of Kigali against differents insecticies such as permethrin, alphacypermethrin and deltamethrin revealed a resistance intensity. The moratility recorded in Kicukiro district was 94% for alphacypermethrin, at 69% for deltamethrin and at 63% for permethrin (Fig 6).

**Figure 6.**
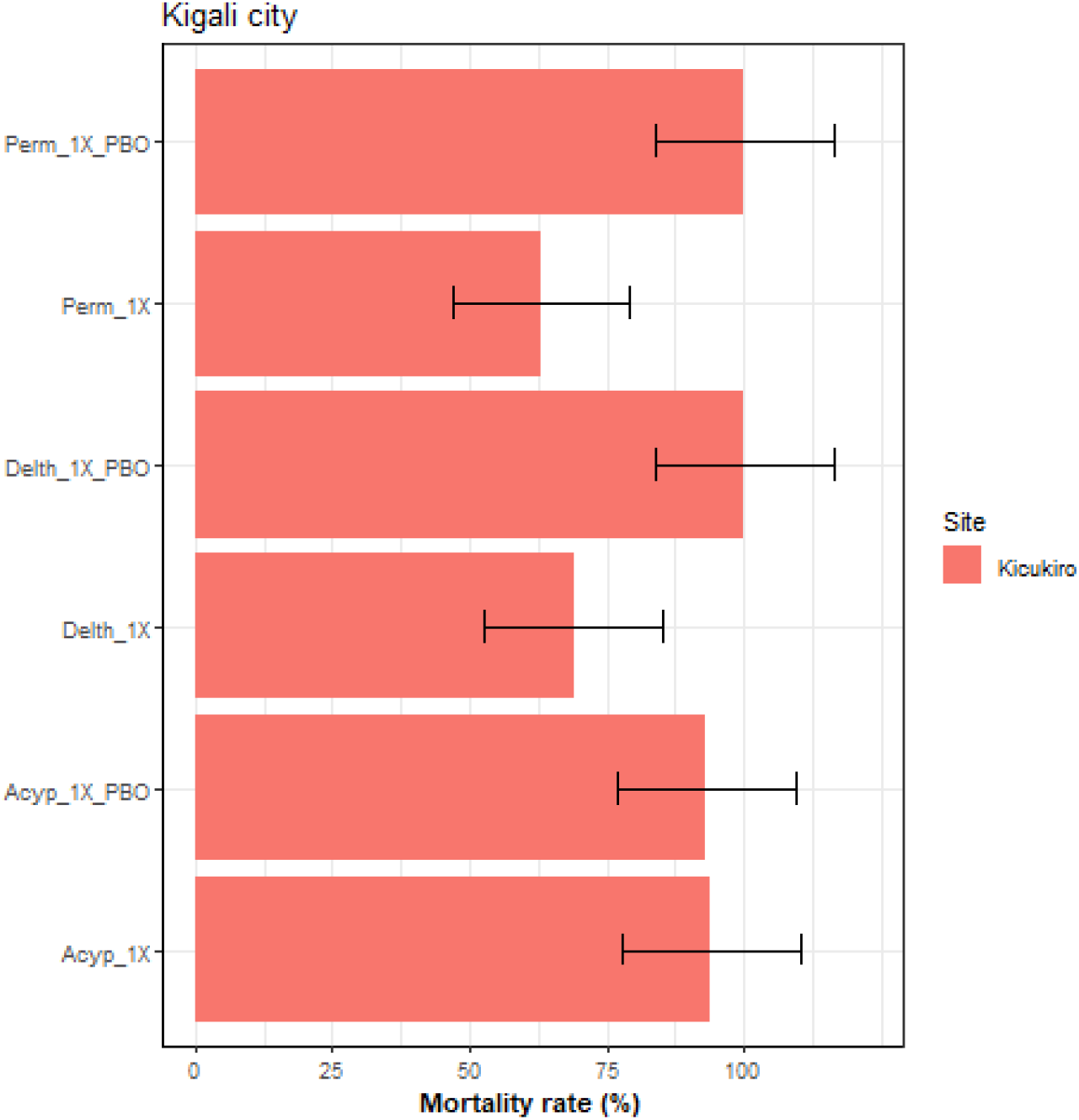
Susceptibility profile of *An. gambiae* population from city of Kigali. *An. gambiae* population were exposure to the three insecticides such as permethrin (Perm_1X), deltamethrin (Delth_1X) and alphacypermethrin (Acyp_1X). The effect of pre-exposure to synergist PBO against pyrethroids also represented (Perm_1X_PBO, Delth_1X_PBO, and Acyp_1X_PBO). Silicon oil was used as a control (mortality ≤5%). Results are average of percentage mortalities and error bars represent ±SE of mean.

### 3.6 Knockdown effect of insecticide

The lowest knockdown rate with alphacypermethrin (Table 1.) was observed in Rwesero (39%) and Gashora (47%), while in permethrin (Table 2), the least was observed in Rwesero and Gashora with 38% and 42% respectively. Although, both Bungwe and Mbuga had the highest knockdown rate of 100% and 97% respectively for permethrin and alphacypermethrin separately. Rukara, Remera and Rwesero represents the lowest knockdown rate with 55% each for deltamethrin (Table 3), while Bungwe and Gashora showed the highest knockdown rate of 100% each.

**Table 2.**
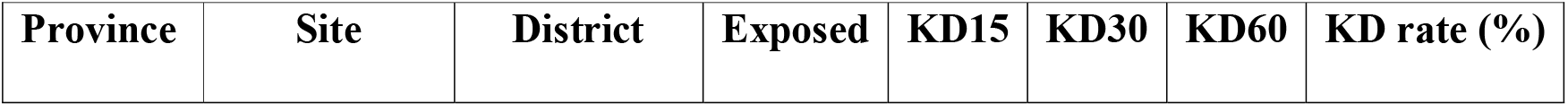
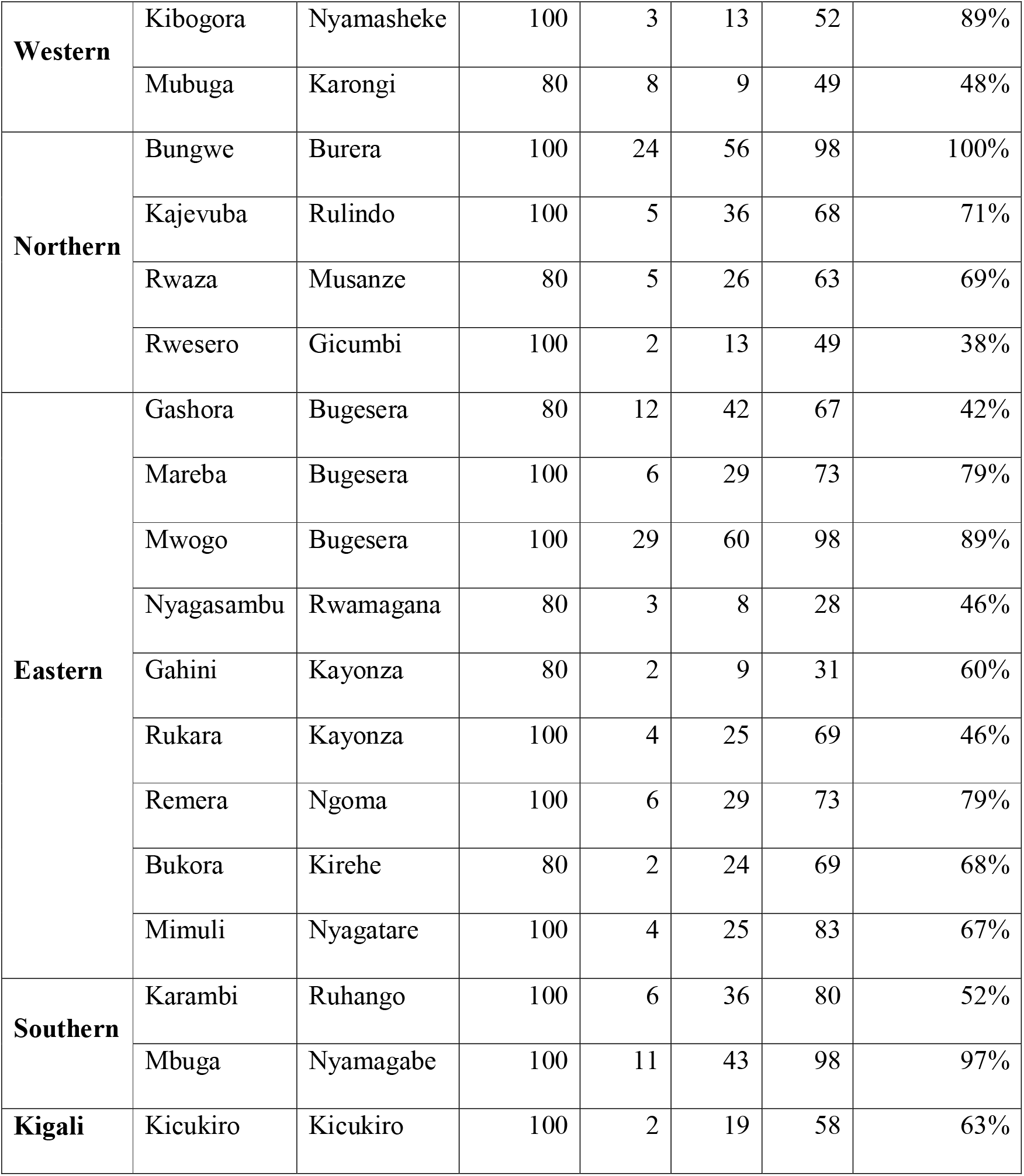
Knockdown effect of permethrin.

**Table 3.** Knockdown effect of deltamethrin.

| **Province** | **Site** | **District** | **Exposed** | **KD15** | **KD30** | **KD60** | **KD rate (%)** |
| --- | --- | --- | --- | --- | --- | --- | --- |
| **Western** | Kibogora | Nyamasheke | 100 | 9 | 51 | 80 | 98% |
|  | Mubuga | Karongi | 80 | 4 | 54 | 72 | 73% |
| **Northern** | Bungwe | Burera | 100 | 20 | 55 | 98 | 100% |
|  | Kajevuba | Rulindo | 100 | 40 | 76 | 99 | 94 |
|  | Rwaza | Musanze | 80 | 16 | 74 | 80 | 78% |
|  | Rwesero | Gicumbi | 100 | 7 | 31 | 90 | 55% |
| **Eastern** | Gashora | Bugesera | 80 | 26 | 64 | 80 | 100% |
|  | Mareba | Bugesera | 100 | 25 | 49 | 76 | 57% |
|  | Mwogo | Bugesera | 100 | 19 | 44 | 81 | 59% |
|  | Nyagasambu | Rwamagana | 80 | 32 | 69 | 80 | 79% |
|  | Gahini | Kayonza | 80 | 16 | 51 | 80 | 78% |
|  | Rukara | Kayonza | 100 | 0 | 33 | 75 | 55% |
|  | Remera | Ngoma | 100 | 11 | 32 | 74 | 55% |
|  | Bukora | Kirehe | 80 | 29 | 53 | 80 | 79% |
|  | Mimuli | Nyagatare | 100 | 1 | 25 | 64 | 66% |
| **Southern** | Karambi | Ruhango | 100 | 48 | 90 | 98 | 97% |
|  | Mbuga | Nyamagabe | 100 | 36 | 81 | 100 | 98% |
| **Kigali** | Kicukiro | Kicukiro | 100 | 5 | 27 | 69 | 69% |

## 4 Discussion

In Rwanda like other African countries uses insecticides for crops and public health. In western province we found a high intensity resistance in mosquitoes collected in Mubuga, Karongi district where the mortality rate was 48% to permethrin, 54% to alphacypermethrin and 73% to deltamethrin respectively (Fig. 2). In Eastern province, the bioassay revealed a resistance to permethrin, alphacypermethrin and deltamethrin on the mosquitoes collected in Mimuli, Mwogo, and Mareba, with mortality rate ranging from 57% to 89%. However, the synergist assay with PBO indicated that metabolic resistance mechanism is fully involved in mosquitoes collected in Nyagatare district (Mimuli), Bugesera district (Mwogo and Mareba) with mortality rate of 100% (Figure 3A). This *An. gambiae* population collected in Nyagasambu, Gashora, Bukora, Gahini was also resistant to permethrin and Alphacypermethrin with a mortality rate observed between 42% to 80%. Subsequently, the use of PBO indicated the mortality rate of 80% in three districts such ad Rwamagana district (Nyagasambu), Bugesera district (Gashora), and Kayonza district (Gahini). Thus, the metabolic mechanism was partially involved (Fig 3A and 3B) while the insecticide resistance carried out to pyrethroids from 2011 to 2013 was detected the gradual increase of resistance in Rwanda [7]. *An. gambiae* population from Bungwe (Burera district) are full susceptible to permethrin, deltamethrin and alphacypermethrin with a mortality rate of 100%. In Northern province, we found that the *An. gambiae* population from Rwesero (Gicumbi district) showed a high resistance to permethrin, alphacypermethrin and deltamethrin with mortality rate of 38%, 39% and 55% respectively. Though, the *An. gambiae* population become fully susceptible after exposure to PBO with mortality rate of 100%, thus the metabolic resistance mechanism is fully involved (Fig. 4). In Southern province, the *An. gambiae* population from Karambi (Ruhango district) was resistance to permethrin and alphacypermethrin which showed a mortality rate of 52% and 72% respectively. Therefore, after exposure to PBO, mosquitoes population become fully susceptible with mortality rate ranging between 98-100%, which indicated that metabolite resistance is fully involved (Fig 5) in that region. The *An. gambiae* collected from Kicukiro (Kicukiro district) located in City of Kigali, showed a resistance to Alphacypermethrin where mortality rate recorded was 94% and after exposure to PBO the mortality observed was 93%, which shed light on role of other resistance mechanism involved rather than metabolic resistance mechanism. On other the *An. gambiae* showed a resistance to permethrin and deltamethrin then become fully susceptible after exposure to PBO (mortality = 100%). This indicates the involvement of metabolic resistance in *An. gambiae* from Kicukiro (Fig 6). Our results line with previous studies, concerning expansion of pyrethroid resistance through agricultural activities [10, 11].

We found that metabolic resistance is likely the major resistance mechanism in mosquitoes population exposed to permethrin collected from various districts (Fig. 7) and per sentinel such as in Nyagatare district (Mimuli), Ngoma district (Remera), Kayonza district (Rukara), Bugesera district (Mareba, Mwogo), Nyamasheke district (Kibogora), Kicukiro district (Kicukiro), Gicumbi district (Rwesero), Ruhango district (Karambi). Most of the insecticide used in Rwanda belong to the similar chemical classes [12], which might force mosquitoes to use metabolic resistance mechanism. The near full recovery of susceptibility was observed to permethrin after exposure to PBO (mortality was between 98% to 100%).

**Figure 7.**
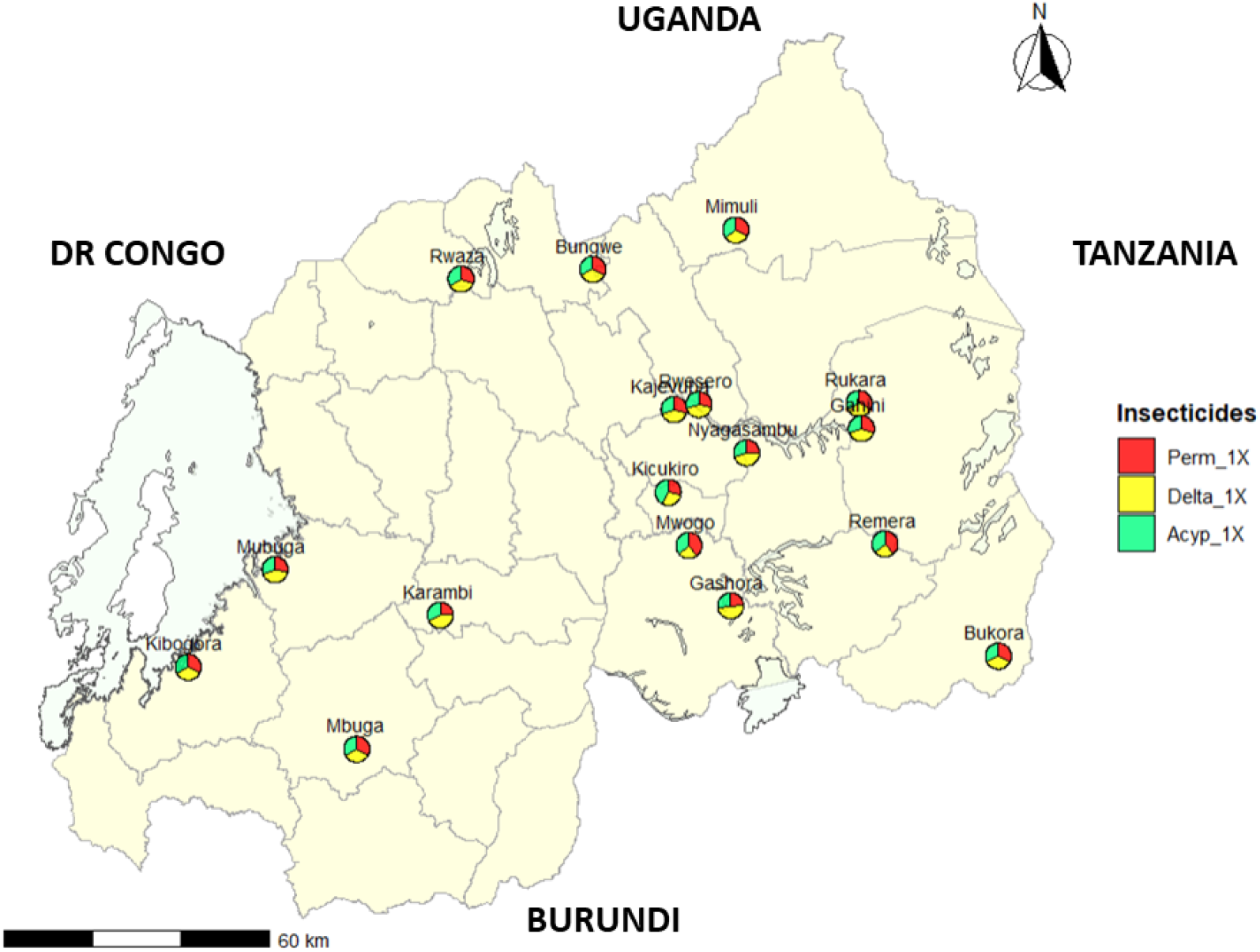
Spatial distribution of insecticide resistance profile in Rwanda. Mosquito population were exposed to the three insecticides such as permethrin (Perm_1X), deltamethrin (Delta_1X) and alphacypermethrin (Acyp_1X). The pie chart represents the mortality rate (%) per insecticide in each study sites.

The susceptibility after PBO exposure was similar reported in Uganda where P450 metabolic genes such as CYP9K1, CYP6P1, CYP6P3, CYP6P4, and CYP6Z1 have been demonstrated to drive resistance to pyrethroids including permethrin, deltamethrin, and alphacypermethrin [13]. Uncertain, metabolic resistance (cytochrome P450s, carboxylesterases and glutathione S-transferases) may be the main pyrethroid resistance in several districts in Rwanda and further investigations have to be performed to elucidate the status of resistance.

## Conclusion

Our study found that the high resistance to pyrethroid insecticides were recorded in *An. gambiae* population from various districts across Rwanda, and showed a partial recovery of susceptibility for pyrethroids using PBO synergist assay. Although further studies are needed to underlying molecular basis of the resistance, whether metabolic resistance, and genetic factors either in mosquito populations or environmental conditions.

## Ethics statement

Permits and consents were not needed because the study was limited to mosquitoes only and did not involve human participants.

## Study limitation

The major limitations of this study are lack of bioassay results of insecticides after PBO pre-exposure in some study sites and not maintain larvae collection over time to evaluate insecticide resistance in the same study sites. These important studies will be the subject of future work.

